# Single-molecule nanopore-tweezers analysis of transcription elongation by intact and “stalk-less” yeast RNA polymerase II

**DOI:** 10.64898/2026.07.20.739707

**Authors:** Shuya Yang, Matthew Chen, Andrew H. Laszlo, Jonathan M. Craig, Jens H. Gundlach, Craig D. Kaplan, Richard H. Ebright

## Abstract

<u>S</u>ingle-molecule <u>p</u>icometer-<u>r</u>esolution <u>n</u>anopore tweezers (SPRNT) enables monitoring of translocation of a nucleic-acid motor protein on a nucleic-acid track with sequence registration, sub-nucleotide spatial resolution, sub-millisecond temporal resolution, and the ability to apply forces that assist or oppose translocation. Recently, we used SPRNT to analyze the translocation of single molecules of *Escherichia coli* RNA polymerase relative to the DNA template strand during transcription elongation, and we directly detected sequence-dependent pausing and formation of a “half-translocated state” at the *E. coli yrbL* consensus pause element. Here, we apply SPRNT to analyze the translocation of single molecules of yeast RNA polymerase II (Pol II) relative to the DNA template strand during transcription elongation with single-nucleotide spatial resolution and millisecond-scale temporal resolution at biologically relevant, saturating substrate concentrations; we compare translocation by intact, 12-subunit Pol II to translocation by a 10-subunit Pol II sub-assembly lacking the dissociable Rpb4-Rpb7 Pol II “stalk”; and we assess possible pausing by intact Pol II and stalk-less Pol II at the *E. coli yrbL* consensus pause element. The results show that intact Pol II elongates more rapidly than stalk-less Pol II and show that neither intact Pol II nor stalk-less Pol II pauses at the *E. coli yrbL* consensus pause element.

**One-sentence summary:** Nanopore tweezers enable monitoring of translocation of RNA polymerase II relative to DNA in transcription elongation with single-nucleotide spatial resolution and millisecond-scale temporal resolution at biologically relevant, saturating substrate concentrations.

## Introduction

RNA polymerase II (Pol II) is the enzyme that performs mRNA synthesis in eukaryotes (1–2). Pol II comprises 12 subunits (Rpb1-Rpb12; “intact Pol II”; Fig. 1A) and can be dissociated to yield a catalytically active core sub-assembly comprising ten subunits (Rpb1-Rpb3, Rpb5-Rpb6, and Rpb8-Rpb12; “stalk-less Pol II”; Fig. 1B) and a “stalk” sub-assembly comprising two subunits (Rpb4 and Rpb7) (1–5). Biochemical and genetic results indicate that the Pol II stalk is essential for transcription initiation and plays roles in regulation of transcription but is not essential for transcription elongation (1–5).

**Figure 1.**
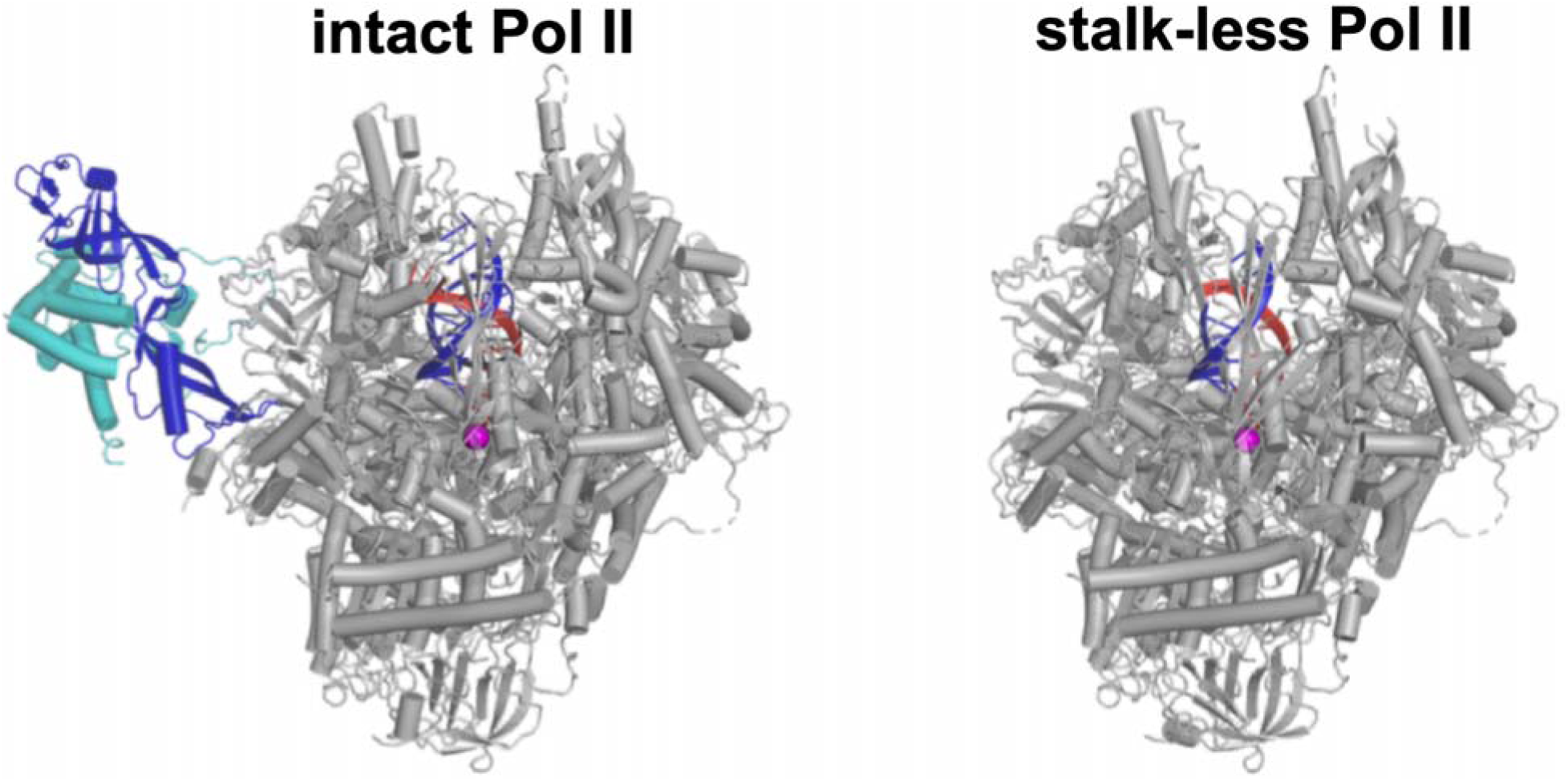
Intact Pol II and stalk-less Pol II. Structures of TECs containing intact Pol II (left; PDB 1Y1W; 47) and stalk-less Pol II (right; PDB 1I6H; 48). Gray, Pol II catalytic core comprising subunits Rpb1-Rpb3, Rpb5-Rpb6, and Rpb8-Rpb12; cyan and blue, Pol II stalk, comprising subunits Rpb4 and Rpb7; violet sphere, Pol II active-center catalytic Mg^2+^ ion; black, blue, and red ribbons, nontemplate DNA strand, template DNA strand, and RNA.

During transcription elongation, an RNA polymerase functions as a molecular motor that translocates relative to the DNA template strand as it synthesizes an RNA copy of the DNA template strand. Translocation of single molecules of *E. coli* RNA polymerase relative to DNA during transcription elongation has been analyzed with accurate sequence registration and single-nucleotide resolution by use of optical tweezers (6–7). However, limitations on the spatiotemporal resolution of optical tweezers have restricted analysis with accurate sequence registration and single-nucleotide resolution to studies that employ nucleoside triphosphate (NTP) concentrations multiple orders of magnitude lower than those in cells and that involve nucleotide-addition rates an order of magnitude lower than those in cells, precluding analysis of biologically relevant reaction conditions and kinetics (6–7).

Recently, we demonstrated the application of <u>s</u>ingle-molecule <u>p</u>icometer-<u>r</u>esolution <u>n</u>anopore tweezers (SPRNT; 8-13) to monitor the translocation of single molecules of *Escherichia coli* RNA polymerase relative to the DNA template strand during transcription elongation with accurate sequence registration and unprecedented spatiotemporal resolution, enabling analysis of biologically relevant reaction conditions and kinetics (14), and we directly detected sequence-dependent transcriptional pausing and formation of a transcriptionally inactive state with fractional-nucleotide translocation--”half-translocated state”--at the *E. coli yrbL* consensus pause element (14).

Here, we demonstrate the application of SPRNT (8–14) to monitor the translocation of single molecules of yeast Pol II relative to the DNA template strand during transcription elongation with accurate sequence registration and unprecedented spatiotemporal resolution, enabling analysis of biologically relevant reaction conditions and kinetics; we analyze translocational behaviors of intact Pol II and stalk-less Pol II at pause-free sequences; and we analyze translocational behaviors of intact Pol II and stalk-less Pol II at the *E. coli yrbL* consensus pause element.

## Methods

### Yeast Pol II and yeast stalk-less Pol II

Biotinylated yeast intact Pol II and biotinylated yeast stalk-less Pol II were prepared as described (15) and were stored in aliquots in 20 mM Tris (pH 8.0), 40 mM KCl, 5 mM MgCl_2_, 1 mM dithiothreitol, and 10% glycerol at -80°C.

### Nucleic-acid scaffold

SPRNT was performed using the nucleic-acid scaffold of Figs. 2A and S1. The nucleic-acid scaffold comprises the T7 A1 +27 force-dependent transcriptional arrest site (14; 16) followed by a transcription unit containing the *E. coli yrbL* bacterial consensus pause element (14; 17) and is identical to the nucleic-acid scaffold “Sequence 1” used in SPRNT analysis of transcription elongation by *E. coli* RNA polymerase (14), except that the cholesterol “membrane anchor” (10–14) is incorporated by annealing of a cholesterol-TEG-containing “adaptor” oligodeoxyribonucleotide to the 3’-end of the nontemplate strand, rather than being covalently incorporated at the 3’-end of the nontemplate strand. Oligodeoxyribonucleotides and oligoribonucleotides were purchased (Stanford University Protein and Nucleic Acid Facility), PAGE-purified, dissolved in 10 mM Tris-Cl, pH 8.5, to 100 μM, and stored at -20°C in aliquots. The nucleic-acid scaffold was prepared by mixing 0.5 µM nontemplate-strand oligodeoxyribonucleotide (5’-GAATAGCCATCCCTATCATCGACATTGCACAACTGCGCCAGCTACTAGC ACCCTTTTTTTTTTTTTTTTTTCAGGCTCAGTACGATCAGTATCC -3’), 0.5 µM template-strand oligodeoxyribonucleotide (5’-GGGTGCTAGTAGCTGGCGCAGTTGTGCAATGTCGATGATAGGGATGGCTATTCGCCGTGTCCCGACTCCTCATGCAGGTCGTTTTTTTTTTTTTTTTTTTTTTTTT-3’), 0.6 µM oligoribonucleotide (5’-rCrGrUrUrArGrArGrGrGrArCrArCrGrGrCrGrArArUrArGrCrCrArU-3’), and 2.5 μM adaptor oligodeoxyribonucleotide [5’-(cholesterol-TEG)-(T)_50_-(Spacer18)-GGATACTGATACTGAGCCTG-3’] in 100 µl annealing buffer (2 mM HEPES-NaOH, pH. 8.0, 50 mM NaCl, and 1 mM EDTA), heating 3 min at 95°C, and cooling to 20°C in 1°C steps with 1 min per step using a thermal cycler. The resulting nucleic-acid scaffold was aliquotted and stored at -80°C.

**Figure 2.**
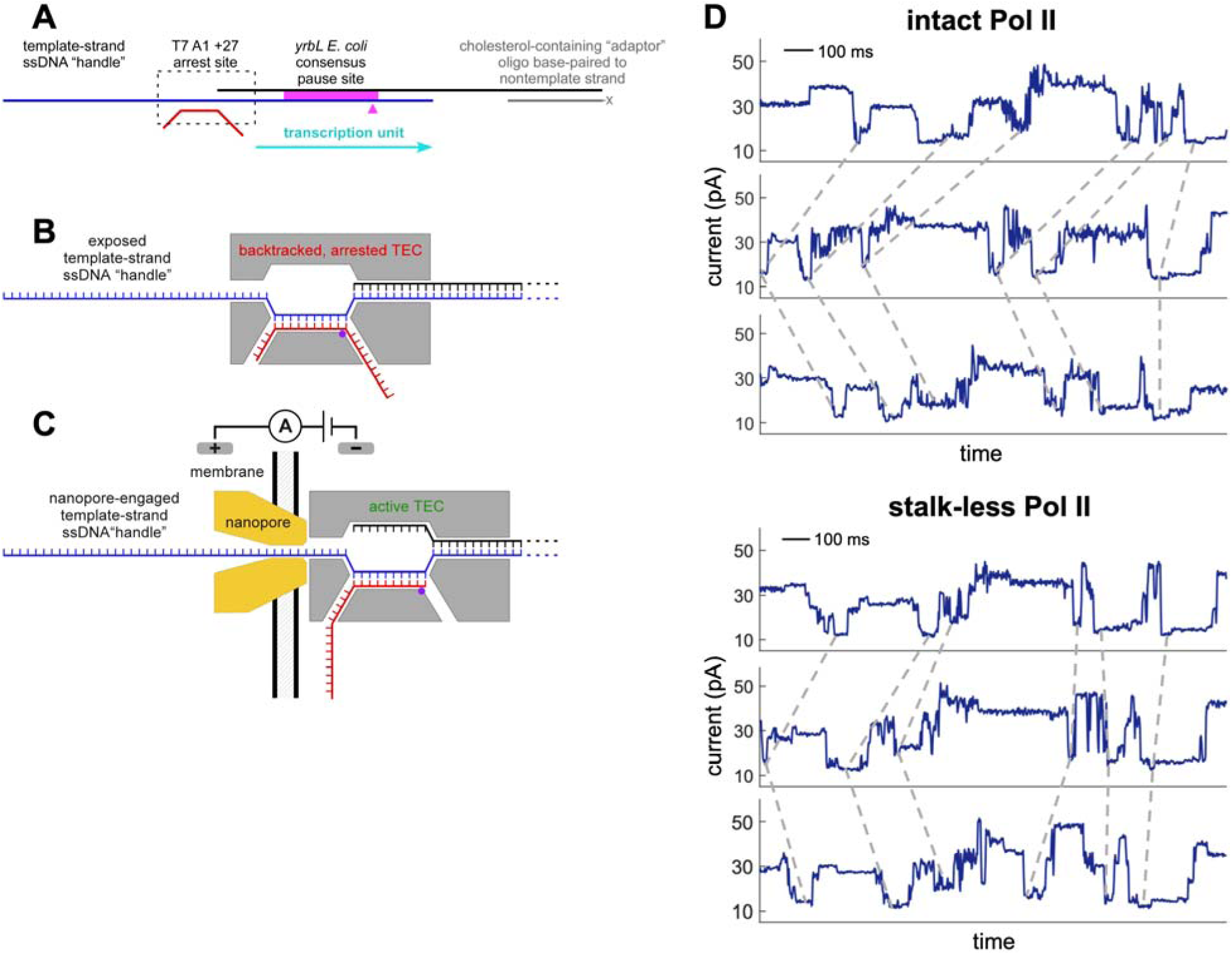
SPRNT analysis of intact Pol II and stalk-less Pol II: experimental design. **(A)** Nucleic-acid scaffold analyzed. Black, blue, red, and gray lines, nontemplate-strand DNA oligonucleotide, template-strand DNA oligonucleotide, RNA oligonucleotide, and cholesterol-containing adaptor oligonucleotide; black dashed rectangle, position at which RNA polymerase binds to a yield back-tracked, arrested TEC at the T7 A1 +27 force-dependent arrest site (14; 16); cyan arrow, transcription unit transcribed after the back-tracked, arrested TEC engages a nanopore, is subjected to voltage-dependent assisting force, and is converted to an active TEC (14); magenta rectangle and magenta arrowhead, *E. coli yrbL* consensus pause element and position within the pause element at which pausing by *E. coli* RNA polymerase occurs (14; 17); “-x,” 5’-cholesterol-TEG-(T)_50_-Spacer18-. (14). **(B)** Backtracked, arrested TEC assembled on synthetic nucleic-acid scaffold containing a T7 A1+27 force-dependent arrest site and containing a ssDNA “handle” that enables capture by a nanopore (14). **(C)** Active TEC formed after a back-tracked, arrested TEC is captured by a nanopore and subjected to voltage-dependent assisting force, triggering RNA polymerase forward translocation, arrest escape, and transcription elongation (14). During transcription elongation, RNA polymerase drives the DNA template strand through the nanopore, and, at each time point, current across the nanopore reports the identity of the template-strand nucleotide in the nanopore constriction (14). **(D)** Representative single-molecule current-vs.-time traces for intact Pol II (top; three current-vs.-time traces) and stalk-less Pol II (bottom; three current-vs.-time traces). Data are from SPRNT reactions at saturating NTPs at 180 mV (∼36 pN assisting force). Black dashed lines indicate selected shared current features.

### Backtracked, arrested TECs

Backtracked, arrested TECs were assembled by adding 10 μl 1 μM intact Pol II and 2 μl 0.5 μM nucleic-acid scaffold in 8 μl transcription buffer (10 mM HEPES, pH 8.0), and 100 mM KCl, 10 mM MgCl_2_, and 1 mM dithiothreitol) or by adding 2 μl 5 μM stalk-less Pol II and 2 μl 0.5 μM nucleic-acid scaffold in 16 μl transcription buffer (Fig. 2B).

### SPRNT data collection

Puck assembly, chamber set up, lipid-bilayer formation, and nanopore insertion were performed as described for SPRNT data collection with “backward” MspA nanopores (14). Transcription reactions were performed by adding 0.5 μl 0.05 μM backtracked, arrested TEC in 20 μl transcription buffer to the “*cis*” well, containing 50-100 μl transcription buffer and 1 mM ATP, 1 mM CTP, 1 mM GTP, and 1 mM UTP at 22°C (for reactions with saturating NTPs) or 50-100 μl transcription buffer and 0.01 mM of one NTP and 1 mM of each other NTP at 22°C (for reactions with one limiting NTP), of a chamber at a holding voltage of 36 mV; incubating 5 min at 22°C; applying a driving voltage across the nanopore; and monitoring ion-current flow across the nanopore for 0.5-3 h at 22°C (Fig. 2C). Driving voltages applied were 120 mV, 150 mV, 180 mV, or 200 mV; these driving voltages result in assisting forces of ∼24 pN, ∼30 pN, ∼36 pN, and ∼40 pN, respectively (12–14). Reactions with saturating NTPs were performed using 1 mM ATP, 1 mM CTP, 1 mM GTP, and 1 mM UTP (Figs. 2D and 4-6). Calibration reactions with one limiting NTP and three saturating NTPs were performed using 0.01 mM limiting NTP and 1 mM of each other NTP (Fig. 3A). At least three reactions, using different samples, were performed for each condition. Data were collected at 5,000 Hz and were down-sampled by averaging to 1,000 Hz, yielding a final temporal resolution of 1 ms.

**Figure 3.**
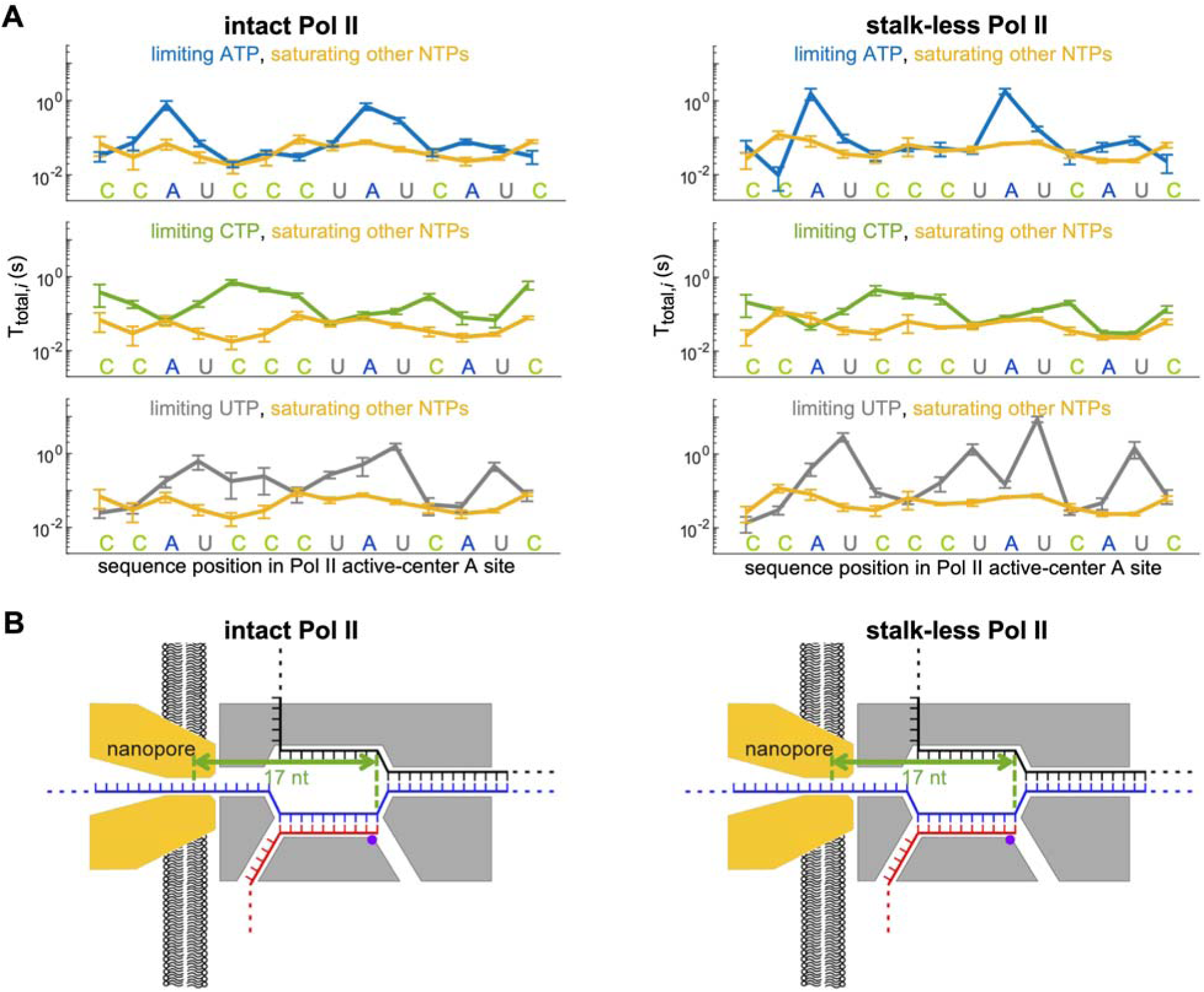
SPRNT analysis of intact Pol II and stalk-less Pol II: determination of “registration distance” (number of nucleotides between template-strand nucleotide in nanopore constriction and template-strand nucleotide in RNA polymerase active-center A site). **(A)** Position-specific dwell times for intact Pol II (left) and stalk-less Pol II (right) from SPRNT reactions with one limiting NTP and three saturating NTPs (i.e., reactions with limiting ATP and saturating other NTPs, reactions with limiting CTP and saturating other NTPs, and reactions with limiting UTP and saturating other NTPs) at 180 mV (∼36 pN assisting force; mean±SEM for 22-60 single-molecule traces). SPRNT reactions with one limiting NTP and three saturating NTPs show long position-specific dwell times at positions where the limiting NTP is the required NTP (i.e., at positions where the limiting NTP is complementary to the template-strand nucleotide in the RNA polymerase active-center A site). Sequence positions are expressed as the required NTP (i.e., as the NTP complementary to the template-strand nucleotide in the RNA polymerase active-center A site). **(B)** Registration distances for intact Pol II (left; 17 nt) and for stalk-less Pol II (right; 17 nt), determined from SPRNT reactions with one limiting NTP and three saturating NTPs by comparing, at each time point, the identity of the template-strand nucleotide in the nanopore constriction (reported by current levels) to the identity of the template-strand nucleotide in the RNA polymerase active-center A site (reported by patterns of long vs. short dwell times).

### SPRNT data analysis

Data pre-processing, current-level finding, current-level-to-position alignment, and preparation of position-vs.-time traces were performed as described (14).

For analysis of pause-free transcription elongation, a position window of position - 0.5 nt to position + 0.5 nt was used, the total dwell time at each position *i* for each TEC *j* was designated as T_total,*ij*_, the mean total dwell time at each position *i* for all TECs was designated as mean T_total,*i*_, the mean total dwell time at all positions for each TEC *j* was designated mean T_total,*j*_, the mean of mean T_total,*i*_ values for all positions was designated as mean T_total_ (averaged over positions), the mean of mean T_total,*j*_ values for all TECs was designated as mean T_total_ (averaged over TECs), and mean elongation velocities were calculated as the reciprocal of mean T_total_ (averaged over positions) or the reciprocal of mean T_total_ (averaged over TECs).

Force-velocity relationships were fitted to the linear Brownian ratchet model described previously for Pol II (18–19):

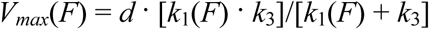

where *Vmax*(*F*) is the elongation velocity at saturating NTPs at force *F*; *k*_1_(*F*) is the rate constant for RNA polymerase forward translocation at assisting force *F*; *k*3 is the combined rate constant for all reactions of the nucleotide-addition cycle after RNA polymerase translocation (i.e., NTP binding, phosphodiester-bond formation, and pyrophosphate release); and *d* is the step size, set at 1 nt; and:

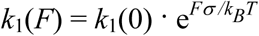

where *k*_1_(0) is the rate constant for RNA polymerase forward translocation at assisting force 0; *F* is the assisting force in N; *σ* is the distance from the energy minimum to the transition state, set at 3.4 10^-10^ m (1 bp; 6), *k_B_* is the Boltzmann constant, and *T* is the temperature in degrees Kelvin.

## Results

### SPRNT analysis of intact Pol II and stalk-less Pol II

We performed SPRNT analysis of intact Pol II and stalk-less Pol II using a synthetic nucleic-acid scaffold substantially identical to the synthetic nucleic-acid scaffold we previously used for SPRNT analysis of *E. coli* RNA polymerase (14; Figs. 2A and S1). The nucleic-acid scaffold contained: (i) the T7 A1 +27 force-dependent transcriptional arrest site (14; 16) followed by a transcriptional unit comprising a first pause-free segment, the *E. coli yrbL* consensus pause element (14; 17), and a second pause-free segment; (ii) a ssDNA “handle” at the 3’ end of the template DNA strand to enable nanopore engagement (8–14); and (iii) a cholesterol “membrane anchor” at the 3’ end of the non-template strand to promote nanopore engagement through interaction with the membrane surrounding the nanopore (10–14) (Figs. 2A and S1).

We collected SPRNT data for intact Pol II and stalk-less Pol II using procedures analogous to the procedures we previously used for SPRNT analysis of *E. coli* RNA polymerase (Figs. 2B-D; and 3; 14). We assembled backtracked, arrested transcription elongation complexes (TECs) by adding of intact Pol II or stalk-less Pol II to the synthetic nucleic-acid scaffold (Fig. 2B). We then added the backtracked, arrested TECs to the “*cis*” compartment of a chamber consisting of a “*cis*” compartment containing transcription buffer and saturating NTPs, a membrane-bound MspA nanopore, and a “trans” compartment; we applied voltage across the nanopore; and we monitored ion-current flow across the nanopore (Fig. 2C). The application of voltage across the nanopore resulted in up to tens of successive reaction cycles, each comprising (i) capture by the nanopore of the exposed template-strand ssDNA “handle” of a backtracked, arrested TEC (Fig. 2B); and (ii) application of assisting force to the backtracked, arrested TEC, triggering forward translocation by the TEC, arrest escape, and active transcription elongation to the end of the transcription unit (Fig. 2C). During each reaction cycle, active transcription elongation by a single TEC drives template-strand DNA through the nanopore over time, yielding current-vs.-time traces (Fig. 2D) that--after data analysis comprising current-level finding, current-to-position alignment, and registration-distance correction--yield position-vs.-time traces that reveal, for each TEC, at each time point, the identity of the template-strand nucleotide in the RNA polymerase active-center addition site (A site) (Figs. 3-4).

**Figure 4.**
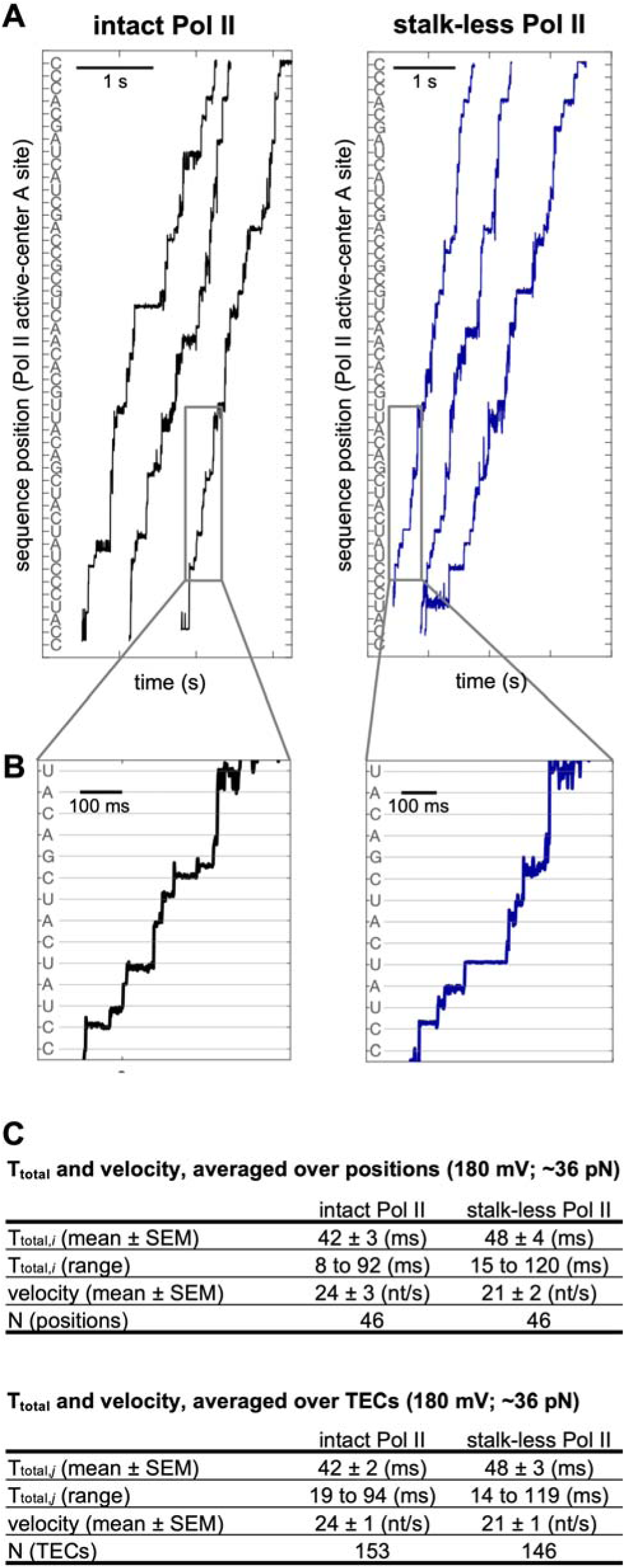
SPRNT analysis of intact Pol II and stalk-less Pol II: single-molecule position-vs.-time traces. **(A)** Representative single-molecule position-vs.-time traces for intact Pol II (left) and stalk-less Pol II (right) from SPRNT reactions at saturating NTPs at 180 mV (∼36 pN assisting force). Sequence positions are expressed as the required NTP (i.e., as the NTP complementary to the template-strand nucleotide in the RNA polymerase active-center A site). **(B)** Enlarged views of representative single-molecule position-vs.-time traces for intact Pol II (left) and stalk-less Pol II (right), showing single-base-pair stepping by (steps at most positions), back-stepping at certain positions (e.g., step from 13th to 14th position from bottom), and possible fractional-base-pair stepping at certain positions (e.g., step from 9th to 10th position from bottom). **(C)** Mean dwell times and elongation velocities for intact Pol II and stalk-less Pol II--averaged over positions (top) or averaged over TECs (bottom)--from SPRNT reactions at saturating NTPs at 180 mV (∼36 pN assisting force).

We determined the registration distance--the number of nucleotides between the template-strand nucleotide in the nanopore constriction and the template-strand nucleotide in the RNA polymerase active-center A site--for intact Pol II and for stalk-less Pol II using procedures analogous to the procedures we previously used to determine the registration distance for *E. coli* RNA polymerase (Fig. 3; 14). To determine the registration distances for intact Pol II and for stalk-less Pol II, we collected and analyzed SPRNT data for reactions in the presence of one limiting NTP and three saturating NTPs (i.e., reactions with limiting ATP and saturating other NTPs, reactions with limiting CTP and saturating other NTPs, and reactions with limiting UTP and saturating other NTPs; Fig. 3A). SPRNT data for reactions with one limiting NTP and three saturating NTPs show long position-specific dwell times at positions where the limiting NTP is the required NTP (i.e., at positions where the limiting NTP is complementary to the template-strand nucleotide in the RNA polymerase active-center A site; Fig. 3A), and thus enable the registration distance to be inferred by comparing, at each time point, the identity of the template-strand nucleotide in the nanopore constriction (reported by the current level in the current-vs.-time trace) to the identity of the template-strand nucleotide in the RNA polymerase active-center A site (reported by the pattern of long vs. short dwell times in the current-vs.-time trace) (Fig. 3B). We find that resulting registration distances are 17 nt for intact Pol II and 17 nt for stalk-less Pol II (Fig. 3B). These registration distances match the registration distance of 17 nt previously reported for *E. coli* RNA polymerase (14), which is structurally and mechanistically homologous to Pol II (1-2; 20).

We collected SPRNT data using driving voltages of 120 mV, 150 mV, 180 mV, and 200 mV (Figs. 2-6); these driving voltages result in assisting forces of ∼24 pN, ∼30 pN, ∼36 pN, and ∼40 pN, respectively (Figs. 2-6; 12-14).

Figs. 4A and 4B presents representative position-vs.-time traces for single TECs of intact Pol II and stalk-less Pol II at saturating NTP concentrations and an assisting force of ∼36 pN. The position-vs.-time traces define, for each TEC, at each time point, the identity of the template-strand nucleotide in the RNA polymerase active-center A site (Figs. 4A and 4B). The spatiotemporal resolution enables detection of single-base-pair stepping by intact Pol II and stalk-less Pol II (Fig. 4B, steps at most positions), enables detection of back-stepping by intact Pol II and stalk-less Pol II at certain positions (Fig. 4B, step from 13th to 14th position from bottom), and enables detection of possible fractional-base-pair stepping at certain positions (Fig. 4B, step from 9th to 10th position from bottom).

Figs. 4C, 5, and S2-S4 present elongation velocities for intact Pol II and stalk-less Pol II from experiments at saturating NTP concentrations and all tested assisting forces. Fig. 6 presents position-specific dwell times for all positions across the transcription unit for intact Pol II and stalk-less Pol II from experiments at saturating NTP concentrations and all tested assisting forces.

**Figure 5.**
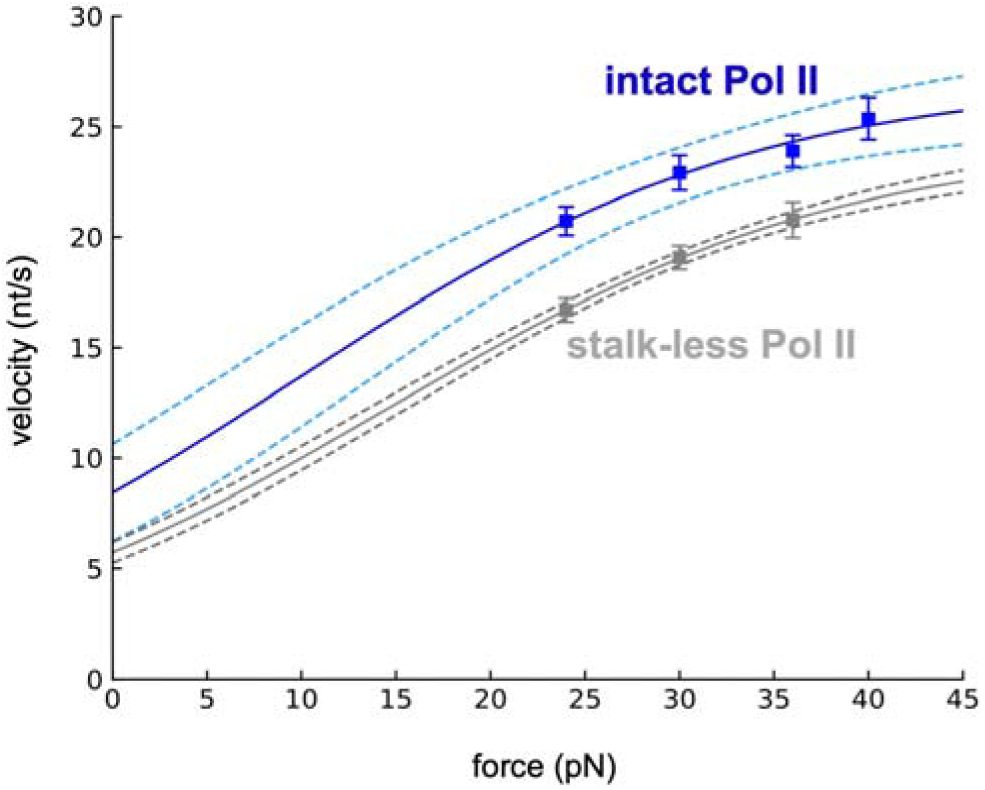
SPRNT analysis of intact Pol II and stalk-less Pol II: elongation velocities and force-velocity relationships. Elongation velocities of intact Pol II (blue symbols) and stalk-less Pol II (gray symbols) from SPRNT reactions with saturating NTPs at 120 mV, 150 mV, 180 mV, and 200 mV (∼24 pN, ∼30 pN, ∼36 pN, and ∼40 pN assisting forces; mean±SEM for 104-167 single-molecule traces) and fits to the linear Brownian ratchet model of refs. 18-19 (blue lines and gray lines; 90% confidence intervals shown as blue dashed lines and gray dashed lines; see Methods for details of fitting). Extrapolated elongation velocities at zero force for intact Pol II (y-intercept of blue line) and stalk-less Pol II (y-intercept of gray line) are 8.4 nt/s and 5.7 nt/s, respectively.

### Elongation velocities of intact Pol II and stalk-less Pol II

Elongation velocities of intact Pol II and stalk-less Pol II at saturating NTPs and all tested assisted forces were force-dependent, increasing with increasing assisted force (Fig. 5), consistent with previous experimental results for Pol II and *E. coli* RNA polymerase (6; 15; 18; 21), and consistent with Brownian-ratchet models that propose translocation by Pol II and *E. coli* RNA polymerase is driven by thermal noise, is biased forward by NTP binding, and is sensitive to assisting forces (6; 15; 18-19; 21).

The observed elongation velocity of intact Pol II is higher than the elongation velocity of stalk-less Pol II at all assisting-force levels tested (Fig. 5). Upon fitting our SPRNT data to the linear Brownian ratchet model (18–19) and extrapolating to zero force, we find that the extrapolated elongation velocity at zero force for intact Pol II is 1.5 times higher than the extrapolated elongation velocity at zero force for stalk-less Pol II (8.4 nt/s vs. 5.7 nt/s; Fig. 5, y-intercepts of blue and gray lines), and the 90%-confidence-interval lower bound of elongation velocity for intact Pol II is higher than 90%-confidence-interval upper bound of elongation velocity for stalk-less Pol II across the entire range of experimental and extrapolated forces (Fig. 5, blue and gray dashed lines). We conclude that the Pol II stalk contributes significantly to the transcription-elongation activity of Pol II. This conclusion is consistent with the structure-based hypothesis that the Pol II stalk may stabilize closed states of the Pol II clamp and thereby increase Pol II processivity (22), with the structure-based hypothesis that RNA binding determinants in the Pol II stalk may interact with RNA emerging from Pol II and thereby increase Pol II processivity (23–24), with the observation that yeast mutants lacking Pol II stalk subunit Rpb4 exhibit growth phenotypes and RNA 3’-end profiles suggestive of decreased Pol II processivity (25–27), and with the observation that the structurally related archaeal RNA polymerase stalk increases archaeal RNA polymerase processivity (28).

We note that the extrapolated elongation velocity at zero force for intact Pol II determined using SPRNT is lower than the elongation velocity at zero force for intact Pol II determined using optical tweezers (8.4 nt/s vs. 20-30 nt/s; Fig 5; 15; 18; 21, 29-30). We attribute this difference to the absence of upstream nontemplate-strand DNA in TECs analyzed using SPRNT (Fig. 2C; 14). Previous results show that the presence of complementary upstream nontemplate-strand DNA in TECs facilitates forward translocation of RNA polymerase by making an additional base pair with template-strand DNA for each base pair of forward translocation (31–33).

### Translocational behaviors of intact Pol II and stalk-less Pol II at *E. coli yrbL* consensus pause element

*E. coli* RNA polymerase engages in sequence-dependent pausing on the time scale of seconds at consensus pause elements (G_-10_Y_-1_G_+1_, where -1 corresponds to the position of the RNA 3′ end and +1 corresponds to the next nucleotide to be incorporated; 14; 17; 34-39). SPRNT results show that *E. coli* RNA polymerase efficiently pauses at the *E. coli yrbL* consensus pause element (17), show that the pause efficiency is >90%, show that the mean pause duration is >1 s, and show that the paused TECs interconvert among five translocational states: (i) backtracked, (ii) pre-translocated, (iii) half-translocated, (iv) post-translocated, and (v) hyper-translocated (14). The SPRNT results further show the half-translocated state is, by far, the most highly occupied state in the paused TECs --accounting for ∼80% of total dwell time at the pause site--is long-lived, and is transcriptionally inactive (14). The results establish that, for *E. coli* RNAP, sequence-dependent pausing at the *E. coli yrbL* consensus pause element predominantly involves a long-lived, transcriptionally inactive, half-translocated state, with half-base-pair translocation of RNAP relative to DNA (14).

It has been unclear whether Pol II recognizes *E. coli* consensus pause elements (36). One group has reported biochemical data indicating pausing by Pol II at an *E. coli* consensus pause element (34), and another group has reported an *in vivo* whole-genome data indicating pausing by Pol II at sequences similar to *E. coli* consensus pause elements (40). However, in both cases, pausing by Pol II occurred at a position adjacent to--not identical to--the position at which pausing by *E. coli* RNA polymerase occurs at an *E. coli* consensus pause element (34; 40); *in vivo* and *ex vivo* whole-genome studies of Pol II pausing other than the study of ref. 34 do not detect pausing by Pol II at sequences similar to *E. coli* consensus pause elements (41–45); and, although not pointed out in the paper reporting the study, an optical-tweezers study of transcription elongation by Pol II on DNA that, by chance, included a sequence matching an *E. coli* consensus pause element, did not detect pausing by Pol II at the sequence (G_-10_T_-1_G_+1_ sequence in Fig. 4, figure supplement 1, of ref. 15).

Here, we used SPRNT to assess pausing by intact Pol II and stalk-less Pol II at an *E. coli* consensus pause element, using exactly the same consensus pause element (*E. coli yrbL* consensus pause element), exactly the same sequences preceding and following the consensus pause element, and exactly the same procedures, as used previously for SPRNT analysis of pausing by *E. coli* RNA polymerase (14; Fig. 6). Our results show that neither intact yeast Pol II nor stalk-less yeast Pol II detectably pauses at the *E. coli* consensus pause element (Fig. 6). Mean position-specific dwell times at the sequence position at which pausing by *E. coli* RNA polymerase occurs, at the adjacent sequence positions, and at all other sequence positions in the consensus pause element were indistinguishable from those in pause-free sequences at all forces tested (26-78 ms, 24-88 ms, and 10-110 ms vs. 7.3-150 ms; Fig. 6). We conclude that neither intact yeast Pol II nor stalk-less yeast Pol II functionally recognizes the *E. coli yrbL* consensus pause element, and we infer that sequence determinants for sequence-dependent pausing differ for Pol II vs. *E. coli* RNA polymerase.

**Figure 6.**
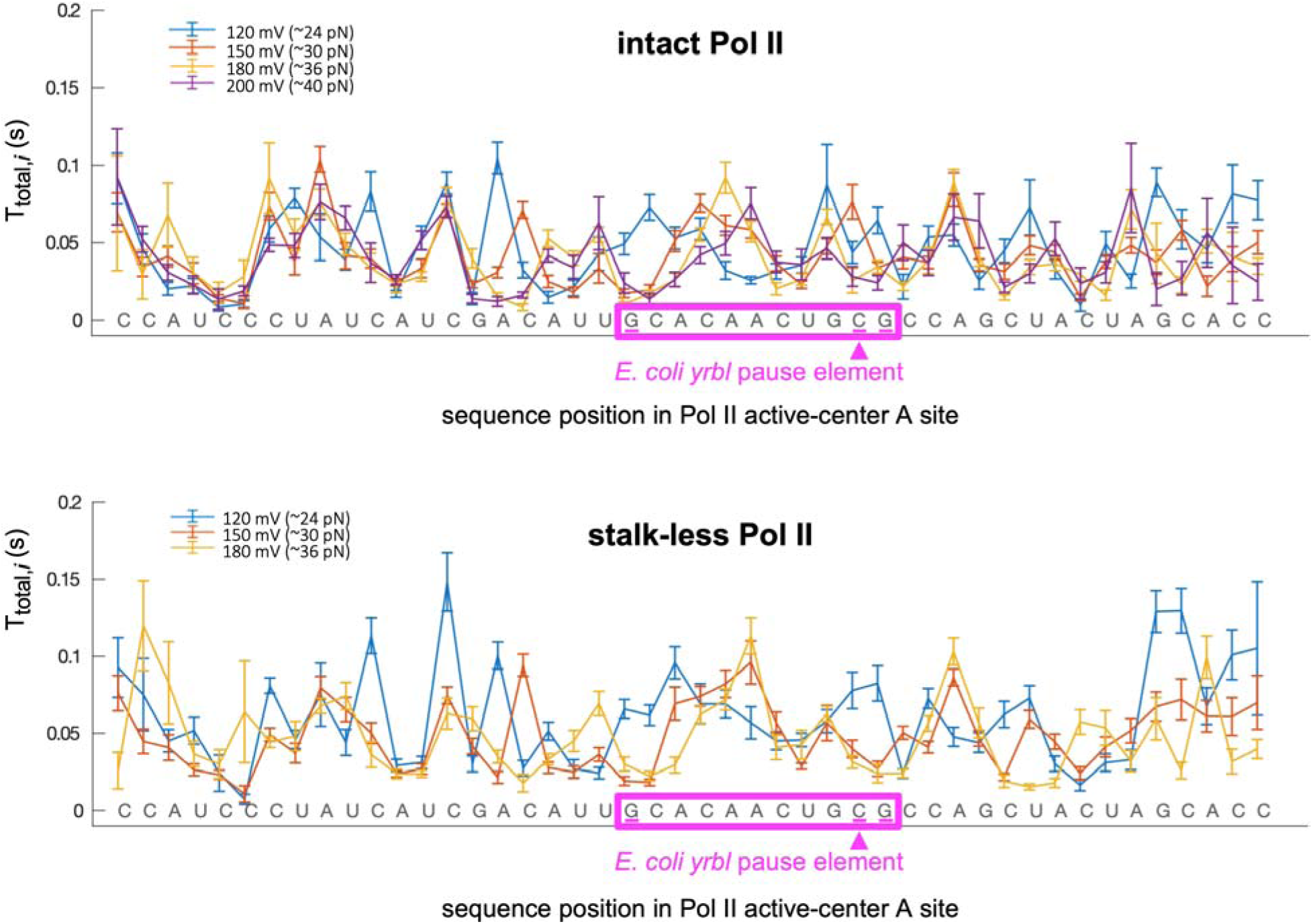
SPRNT analysis of intact Pol II and stalk-less Pol II: position-specific dwell times and absence of pausing at *E. coli yrbL* consensus pause element. Position-specific dwell times for intact Pol II (top) and stalk-less Pol II (bottom) from SPRNT reactions with saturating NTPs at 120 mV (blue), 150 mV (red), 180 mV (yellow), and 200 mV (violet) (∼24 pN, ∼30 pN, ∼36 pN, and ∼40 pN assisting forces; mean±SEM for 104-167 single-molecule traces). Sequence positions are expressed as the required NTP (i.e., as the NTP complementary to the template-strand nucleotide in the RNA polymerase active-center A site). Magenta rectangle and magenta arrowhead, *E. coli yrbL* consensus pause element and position within the pause element at which pausing by *E. coli* RNA polymerase occurs (14; 17).

## Discussion

Our results demonstrate that SPRNT enables detection of single-base-pair stepping by single molecules of Pol II at biologically relevant, saturating NTP concentrations, show that intact Pol II elongates more rapidly than stalk-less Pol II, and show that neither intact Pol II nor stalk-less Pol II recognizes the *E. coli yrbL* consensus pause element. Our results validate the application of SPRNT for single-molecule analysis of transcription elongation, transcriptional pausing, and transcriptional regulation by Pol II.

While this manuscript was in preparation for publication, Vos and co-workers reported deep-sequencing results identifying a Pol II consensus pause element (”consensus super pause”) and reported cryo-EM structural results indicating that Pol II interacts with the consensus super pause in part in a pre-translocated state and in part in a novel “side-tracked” state with single-base-pair back-stepping of Pol II and a previously unobserved standard positioning of the RNA 3’ end (45). The procedures of this report should enable single-molecule analysis of the efficiency, kinetics, and pathway of pausing at the Pol II consensus super pause.

## Acknowledgments

We thank Drs. Ian Nova, Matthew Noakes, Henry Brinkerhoff, and Ian Derrington for training and assistance in SPRNT and Dr. David Degen for assistance in setting up instrumentation for SPRNT at Rutgers University. This work was supported by NIH grants HG005115 to J.H.G., GM097260 and GM144116 to C.D.K., and GM041376 to R.H.E.

## Data Availability Statement

Source data, processed data, and analysis code have been deposited at http://dx.doi.org/10.5281/zenodo.21365436 (46).

## Supplemental Figure Legends

**Figure S1.**
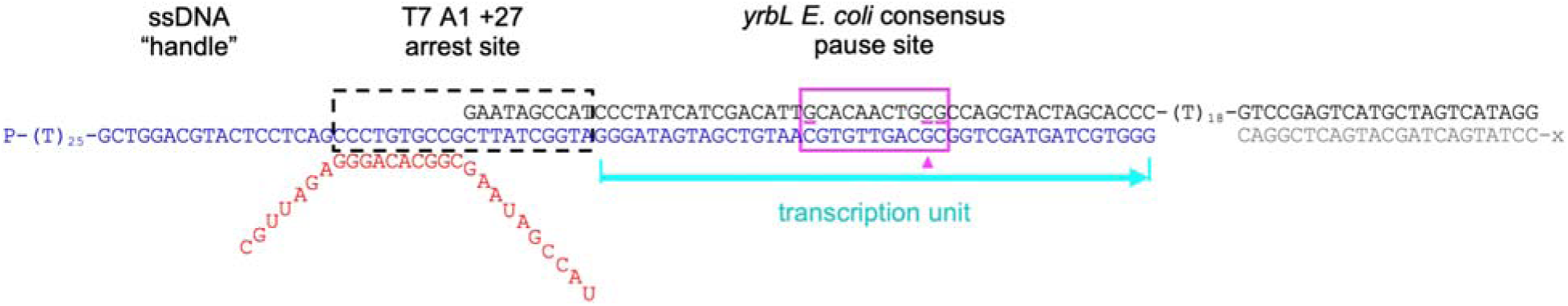
Nucleic-acid scaffold analyzed: sequence. Black, blue, red, and gray sequence, nontemplate-strand DNA oligonucleotide, template-strand DNA oligonucleotide, RNA oligonucleotide, and cholesterol-containing adaptor oligonucleotide; black dashed rectangle, position at which RNA polymerase binds to a yield back-tracked, arrested TEC at the T7 A1 +27 force-dependent arrest site (14; 16); cyan arrow, transcription unit transcribed after the back-tracked, arrested TEC is captured by a nanopore, subjected to voltage-dependent assisting force, and converted to an active TEC (14); magenta rectangle and magenta arrowhead, *E. coli yrbL* consensus pause element and position within the pause element at which pausing by *E. coli* RNA polymerase occurs; “-x,” 5’-cholesterol-TEG-(T)_50_-Spacer18-. (14; 17).

**Figure S2.**
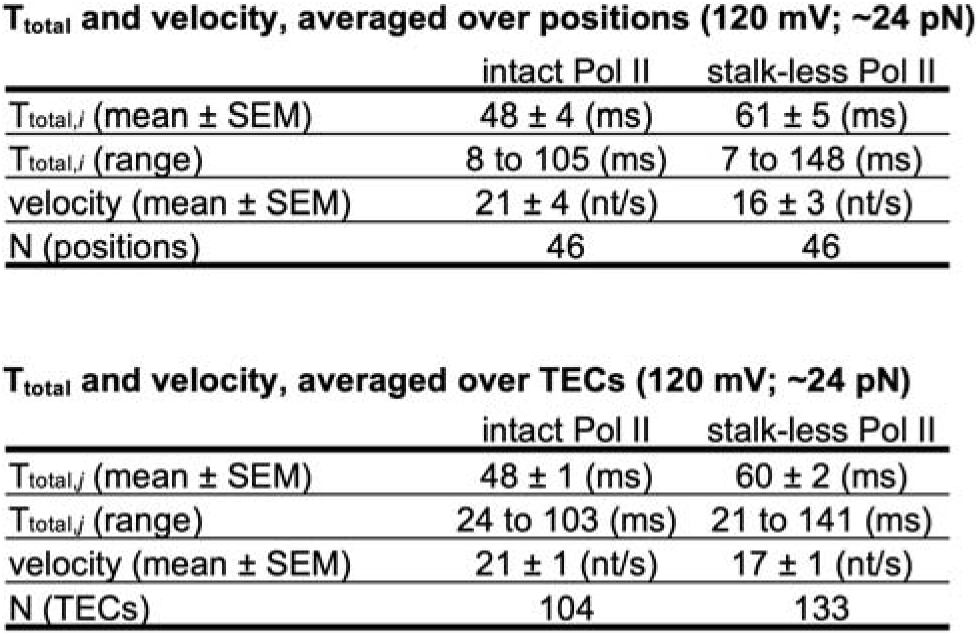
Mean dwell times and elongation velocities for intact Pol II and stalk-less Pol II from SPRNT reactions at saturating NTPs at 120 mV (∼24 pN assisting force).

**Figure S3.**
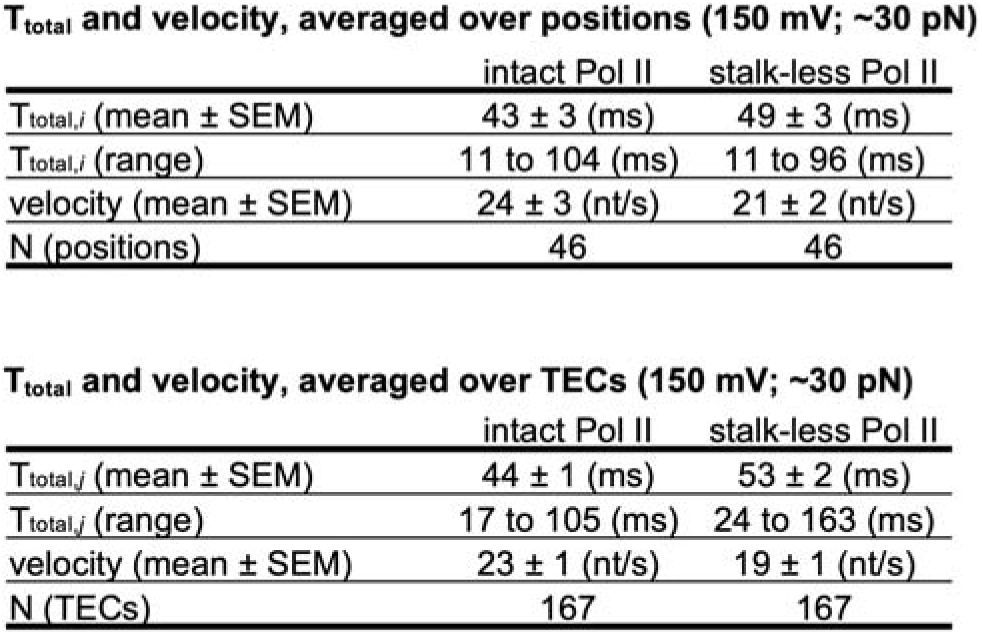
Mean dwell times and elongation velocities for intact Pol II and stalk-less Pol II from SPRNT reactions at saturating NTPs at 150 mV (∼30 pN assisting force).

**Figure S4.**
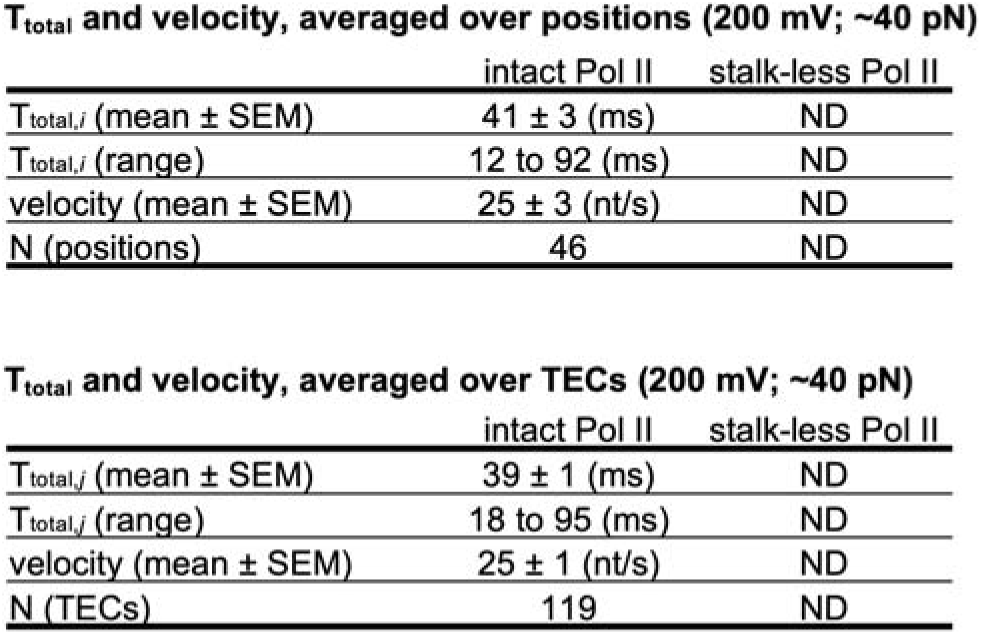
Mean dwell times and elongation velocities for intact Pol II from SPRNT reactions at saturating NTPs at 200 mV (∼40 pN assisting force).

